# Modeling human embryonic adrenogenesis in pluripotent stem cell-derived corticomedullar-like assembloids

**DOI:** 10.64898/2026.09.23.753945

**Authors:** Farah Saleh, Felix James, Dean Rovello, Marcus McIlwain, Andrew Xiao, Caroline Powers, Noah Reichman, Miguel Gonzalez, Barbara Caiazza, Monica Kahn, Zong-Ying Liu, Pushpa Nehru, Carmen Torres, Ting Zhou, Laurie Francoeur, Limor Man, Zev Rosenwaks, Daylon James

## Abstract

During embryogenesis, the cortical and medullary compartments of the adrenal gland emerge from two distinct sources: intermediate mesoderm-derived steroidogenic progenitors that comprise the cortex, and migratory neural crest-derived catecholaminergic progenitors that give rise to the medulla. The distinct cellular origins and paracrine signaling environments that foster induction, differentiation, and/or migration of cortical and medullary layers present challenges to recapitulating their co-emergence and incorporation in a human pluripotent stem cell (hPSC) based system. To address this, we developed an approach for modeling the adreno-gonadal primordium (AGP) in compound organoids that are organized along a gradient of NR5A1 expression and which unexpectedly incorporate SOX10-expressing Schwann cell precursors that ultimately generate chromaffin cells. Deconstruction of paracrine signaling within AGP-like organoids (AGPLOs) identified a fibroblast growth factor 9 (FGF9)-driven mechanism that expands bridge cell-like chromaffin progenitors and correlates to paracrine signaling cues encountered by physiological correlates along their migration corridor to the cortex in vivo. Combination of NR5A1-expressing cells with chromaffin cell clusters generated cortico-medullar-like assembloids (CMLAs) that generated steroids and catecholamines and were responsive to adrenocorticotropic hormone. This work establishes a platform for modeling the reciprocal signaling relationships that drive adrenal gland development and function in health and disease and provides an approach for generating hPSC-derived adrenal cells/tissues that can be applied therapeutically.

## Introduction

The adrenal gland is a critical endocrine organ that regulates metabolism, immune response, blood pressure, and response to stress, and its function is critical for all organ systems. Primary adrenal insufficiency is currently treated with lifelong hormone replacement therapy; however, this does not restore the dynamic adrenal physiology that typifies a healthy axis. In embryonic development, the adrenal gland is generated from two discrete compartments that arise from distinct cellular and anatomic origins^1,2^. The outer adrenal cortex region synthesizes steroid hormones, including cortisol, aldosterone, and adrenal androgens from specialized steroidogenic cells that are organized into three concentric zones; these cells are derived from adrenocortical progenitors of posterior intermediate mesodermal origin that express the transcription factor *NR5A1* (also known as steroidogenic factor 1, SF-1). The inner adrenal medulla region, in contrast, is populated by chromaffin cells that synthesize and secrete catecholamines (epinephrine/norepinephrine) and are neural crest-derived, sharing a common progenitor lineage with sympathetic neurons.

An underappreciated but physiologically critical aspect of adrenal biology is the intimate crosstalk between the cortical and medullary compartments. Adrenal corticosteroids, particularly cortisol, are concentrated within the adrenal medulla owing to direct drainage of cortical sinusoidal blood into the medullary parenchyma^3,4^. This local cortisol gradient induces expression of phenylethanolamine N-methyltransferase (*PNMT*), the enzyme that converts norepinephrine to epinephrine. Therefore, epinephrine, the defining secretory product of the mature adrenal medulla, is produced almost exclusively in adrenal glands that contain a functional cortex^5,6^. Although this cortical-to-medullary signaling axis is inherent to adrenal gland genesis and function, the distinct developmental origins of cortical and medullary cells present significant challenges to generating an integrated adrenal model from human pluripotent stem cells (hPSCs).

While cortical and medullary components of the adrenal gland originate from distinct cell types, these lineages reside within a shared developmental cell field surrounding the adrenogonadal primordium (AGP). The AGP is a transient structure which derives from the coelomic epithelium at embryonic week 4–5 and is populated by *NR5A1*+ progenitors that ultimately give rise to both adrenocortical and gonadal somatic cells^7–9^. The adrenocortical primordium is subsequently invaded by cells originally arising from trunk neural crest cells (NCCs) that form the medullary chromaffin population. Critically, the differentiation and assembly of these distinct progenitor populations are coordinated through paracrine interactions within the cell field that encompasses the AGP. Cortical progenitors provide signals that direct neural crest sympathoadrenal specification^10,11^, while reciprocal neural crest-derived signals modulate cortical progenitor behavior^4^. Platforms employing hPSCs to model organogenesis often fail to reproduce these paracrine co-regulatory signaling relationships, in part because differentiation is directed toward a single target lineage rather than preserving the multi-lineage cellular environment of the primordium itself.

Here, we describe an approach for co-differentiation of cortical and medullary cells in hPSC-derived organoids that recapitulate the AGP cell field. Adreno-gonadal primordium-like organoids (AGPLOs) incorporate cells correlating to AGP-specific derivatives as well as cells sharing the same general cell field within the embryo, including lateral plate mesoderm (LPM), paraxial mesoderm (PM), and NCC derivatives. Paracrine signaling among cell compartments supported downstream differentiation to adrenocortical and chromaffin cells, elucidating a novel fibroblast growth factor 9 (FGF9)-driven mechanism for expanding the sympathoadrenal pool. Control of chromaffin versus adrenocortical fate in AGPLOs enabled combined growth in corticomedullar-like assembloids (CMLAs) that were competent to generate cortex-derived steroid hormones and medulla-derived catecholamines. These findings delineate conditions for selectively tuning differentiation of hPSC-derivatives correlating to the AGP cell field and establish a model of human adrenogenesis that incorporates physiological crosstalk between cortical and medullary compartments.

## Results

### Differentiation of adrenogonadal progenitors in NT^Dim^/NT^Br^ compound organoids

Aiming to reconstitute the cell field in which adrenal progenitors arise from a NR5A1-expressing adrenogonadal primordium, we first generated an hPSC (H9^12^) knock-in reporter line in which the TdTomato fluorescent protein was in incorporated into the NR5A1 locus (**NT** hPSCs, Supp Fig 1a-b). Adapting the differentiation protocol established by Sasaki lab^13^ to support differentiation in adherent conditions (Fig 1a-b), NT**^+^**cells that were FACS-isolated after 16 days of differentiation formed organoids in suspension conditions (Fig 1c-d). After 8 days in culture both NT**^+^** steroidogenic cells expressing STAR, CYP11A1, and CYP17A1 and NT**^neg^** cells of unknown identity (Fig 1e).

**Figure 1.**
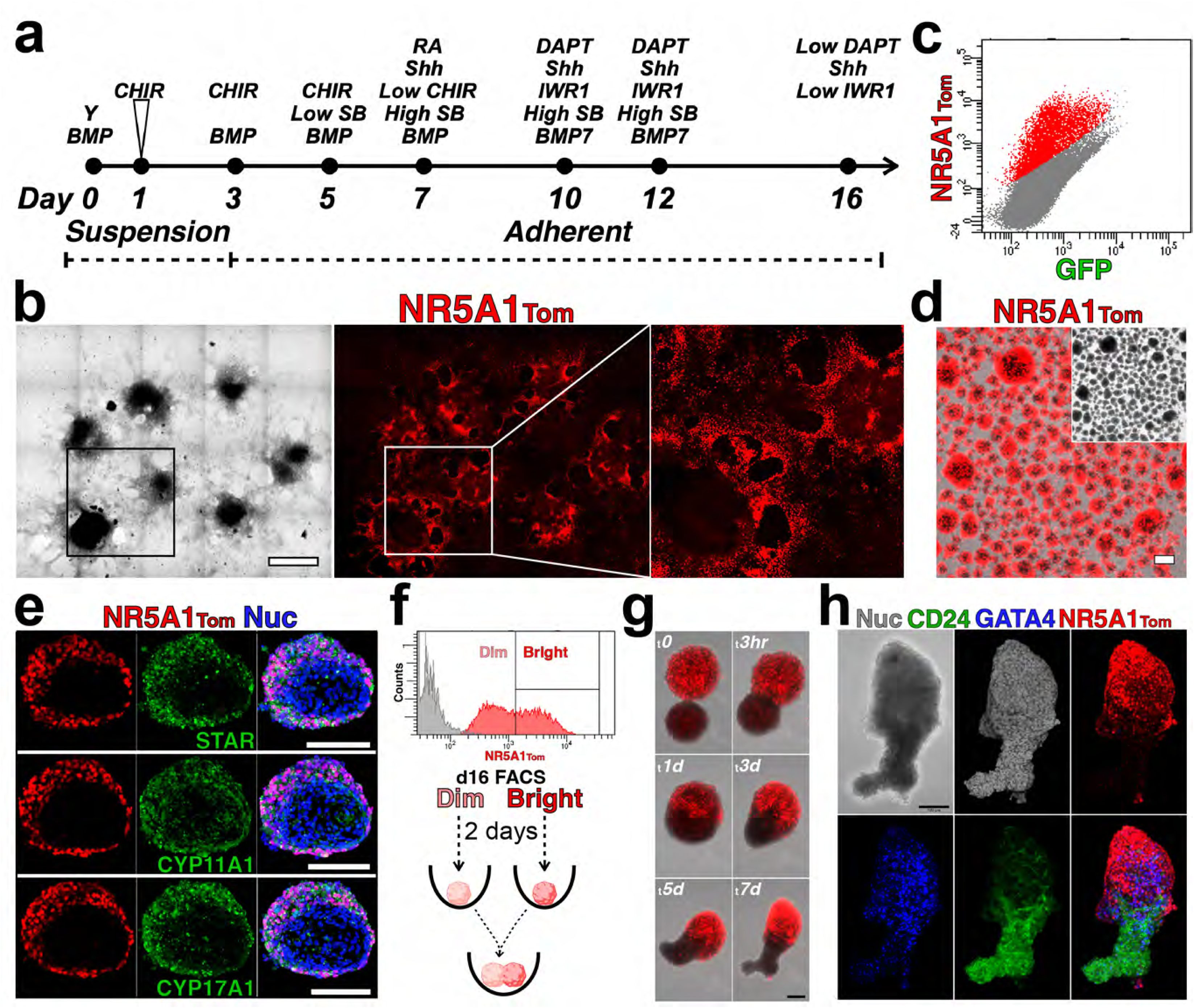
Differentiation of NT hPSCs and generation of AGPLOs. a-b) Differentiation schema for NT hPSCs in suspension and adherent conditions; details provided in Materials and Methods. A wide field view of NT**^+^** colonies is shown in (b). c-d) NT+ cells were sorted in the PE gate (c) and cultured in non-adherent conditions to generate organoids (d). e) Cryosections of organoids after 8 days of culture were immunolabeled with antibodies specific for STAR, CYP11A1, and CYP17A1. f-h) Organoids were generated from NT**^Dim^** and NT**^Br^**sorted cells (f) and combined in compound organoids (AGPLOs) that were subsequently cultured over 7 days with live capture of brightfield and TdTomato channels (g); after terminating the timelapse capture, the AGPLO was recovered and immunolabeled with antibodies specific for CD24 and GATA4. Scale bars – 1 mm in (b), 100 μm in (d, e, g, and h).

Based on the variable expression (∼ 1 log range) of the NT reporter observed among primary differentiation cultures (Fig 1f), we reasoned that this continuum was recapitulating the gradient of NR5A1 expression observed in distinct cell-types of the adrenogonadal primordium as adrenal and gonadal lineages begin to segregate around the 4^th^ week of gestation^9^. To recapitulate the spatial distribution of the adrenogonadal anlage in vitro, we separately generated organoids from cells with high NT expression (NT**^Br^**) and low NT expression (NT**^Dim^**) and recombined them in compound organoids after 2 days (Fig 1g). Timelapse imaging of one representative compound organoid over a week of culture revealed polarization and distribution of cells along a continuum of NT**^Br^** (presumptive adrenal progenitors), GATA binding protein 4 (GATA4**^+^** presumptive gonadal progenitors) and NT**^Negative^**CD24**^+^** (unknown) cells (Fig 1h and Supp Video 1). Together, these results illustrate the capacity for hPSC-derived NT**^Dim/Bright^** cells to reconstitute a field comprised of cells resembling adrenal, gonadal, and other embryonic correlates, hereafter designated as AGPLOs.

### AGPLOs give rise to intermediate mesoderm, paraxial mesoderm, and sympathoadrenal derivatives

To further define the developmental purview of AGPLOs and identify the physiological correlates of undefined cells, we performed single-cell transcriptomic analysis of FACS-isolated NT**^Dim^** (Fig. 1f, ***d0 Dim***) and NT**^Br^** (Fig 1f, ***d0 Br***) cells at 16 days of differentiation, along with AGPLOs that were cultured in a range of recombinant cytokines/small molecules (detailed in Methods) and pooled for analysis after 21 days of differentiation (Fig. 2a, ***d21 Pool***). To provide a physiological basis for comparison, we incorporated previously established single-cell analysis of human fetal adrenogonadal tissues at ∼4 weeks (Fig. 2b, ***Fet AGP***) and ∼5 weeks (Fig 2b, ***Fet Adr***) gestational age^9^. Following quality control and exclusion of poor quality and off-target cells (endothelial, hematopoietic, urogenital epithelium, podocytes, Supp. Fig. 2a-c), remaining populations of AGPLO-derived cells showed significant similarity to primary fetal cells representing chromaffin (PHOX2A, CHGA, SYP), dermomyotome (PAX3, PAX7, MET), adrenocortical (NR5A1, CYP11A1, CYP17A1), adrenogonadal progenitor (STAR), and stromal/mesenchymal fate (TCF21, ACTA2) (Fig 2c-e, and Supp Fig 2d). Additionally lateral plate mesoderm-like progenitors (**LPMProg**, HAND1, WT1) and skeletal muscle-like cells (MYH3, TNNC1) that were not well-represented in primary fetal tissues, as well as rare cells that correlated with Schwann cell precursor (**SCP**, FOXD3, SOX10) identity were identified in AGPLOs (Fig 2d-e and Supp Fig 2d).

**Figure 2.**
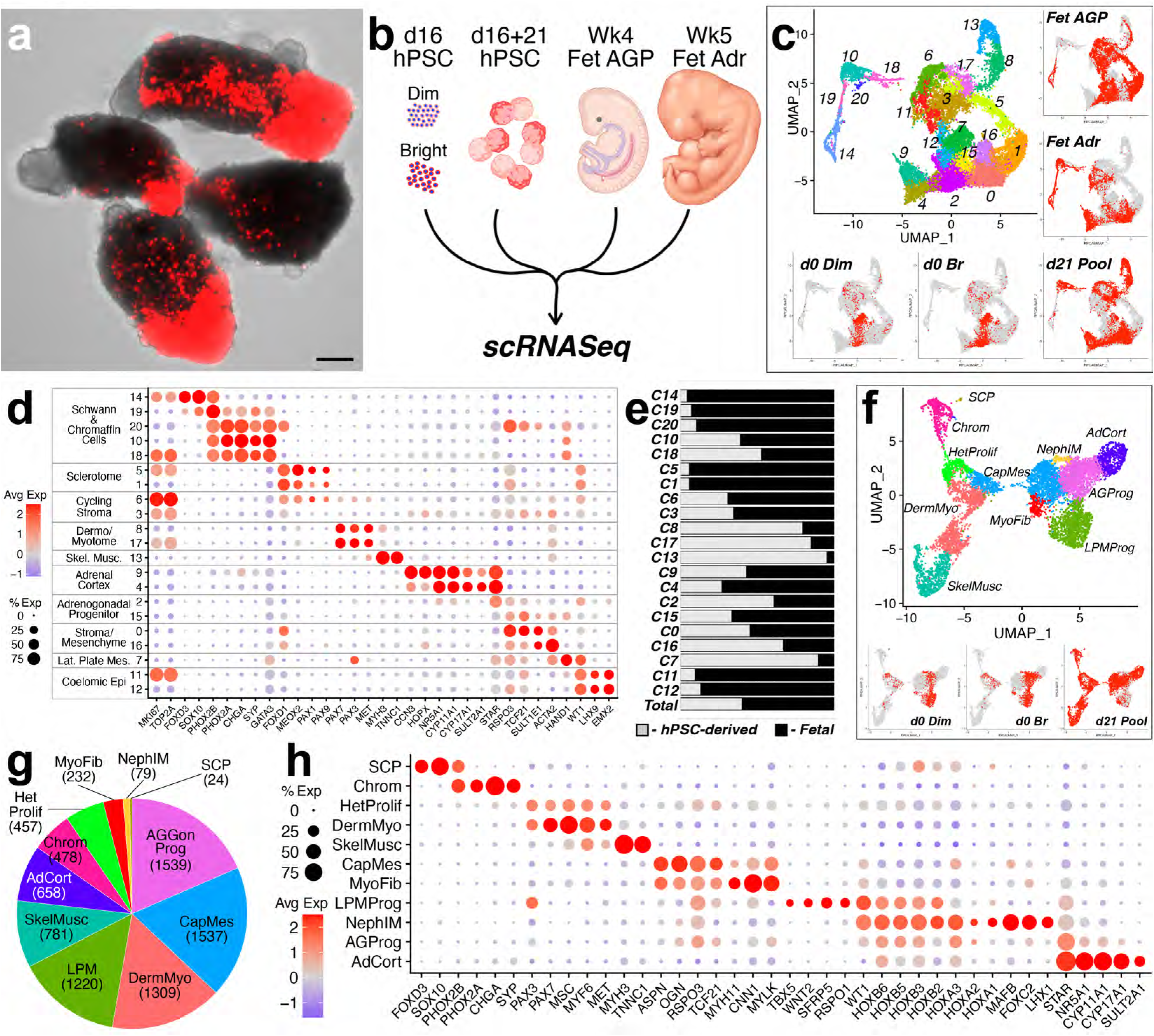
Benchmarking AGPLO-derived populations against fetal correlates. a-b) Following extended differentiation in varied conditions for 21 days (see Materials and Methods), AGPLOs were enzymatically dissociated and, along with freshly isolated NT**^Dim^**and NT**^Br^** sorted cells, processed for single-cell RNA Sequencing and comparison to previously generated (Sasaki lab) fetal libraries (b). c-e) Following quality control and further refinement (see SuppFig 2), overlap of fetal and hPSC-derived cells on the UMAP plot (c), as well as qualitative (d) and quantitative (e) measures of Seurat clusters is shown. f-h) UMAP plots (f) of the hPSC-derived subset from (c), as well as quantitative distribution (g) and transcriptional signatures (h) of annotated clusters. Scale bar in (a) – 100 μm.

In addition to known cellular derivatives of the NR5A1-expressing intermediate mesoderm among the hPSC-derived pool, the presence of **LPMProg** and paraxial mesoderm-derived dermomyotome-like/skeletal muscle-like cells was not unexpected considering the adjacency of these cell types in fetal development. However, the observation of rare SCP-like cells and a significant proportion of chromaffin-like cells (Fig 2c and e) in ***d21 Pool*** indicated incorporation of an ectoderm-derived NCC-like population among the NT**^Dim/Bright^** FACS isolates. Having applied human fetal samples to identify physiological correlates and contextualize the dataset, hPSC-derived cells were isolated, re-clustered, and annotated (Fig 2f-g and Supp Fig 2e-g), demonstrating a variable distribution of cells correlating to sympathoadrenal (**SCP**, chromaffin [**Chrom**]), lateral plate mesodermal (**LPMProg**), intermediate mesodermal (adrenogonadal progenitor [**AGProg**], adrenocortical [**AdCort**], capsular mesenchymal [**CapMes**], myofibroblast [**MyoFib**], nephrogenic progenitor [**NephIM**]), and paraxial mesodermal (skeletal muscle [**SkelMusc**], dermomyotome [**DermMyo**]) fate. Of note, transcriptomic profiling of HOX clusters among the hPSC-derived populations indicated overlap of adrenogonadal derivatives (AGProg, AdCort, CapMes) and SCP within the thoracic zone (Supp Fig 2f-g), corresponding to the axial region where adrenogenesis takes place^14,15^.

### Paracrine signaling within AGPLOs drives expansion of chromaffin cell derivatives

To capitalize on the potential for AGPLOs to recapitulate the paracrine signaling cues that enabled co-differentiation of cells fated to form both mesoderm-derived adrenocortical progenitors and SCP-derived chromaffin/sympathoblast progenitor compartments (Fig. 3a), we examined the expression of canonical ligand/receptor pairs among AGPLO-derived populations (Fig. 3b). Multiple pairings supported the potential for directional signaling between populations involving neurotrophin, fibroblast growth factor (FGF), neuregulin, glial derived neurotrophic factor (GDNF), transforming growth factor beta (TGFb) superfamily, Notch, and Wnt pathways. Additionally, volumetric imaging of AGOs revealed close spatial approximation of chromaffin clusters (SYP**^+^**) and NGFR**^+^** putative SCPs (Fig. 3c and Supp Video 2).

**Figure 3.**
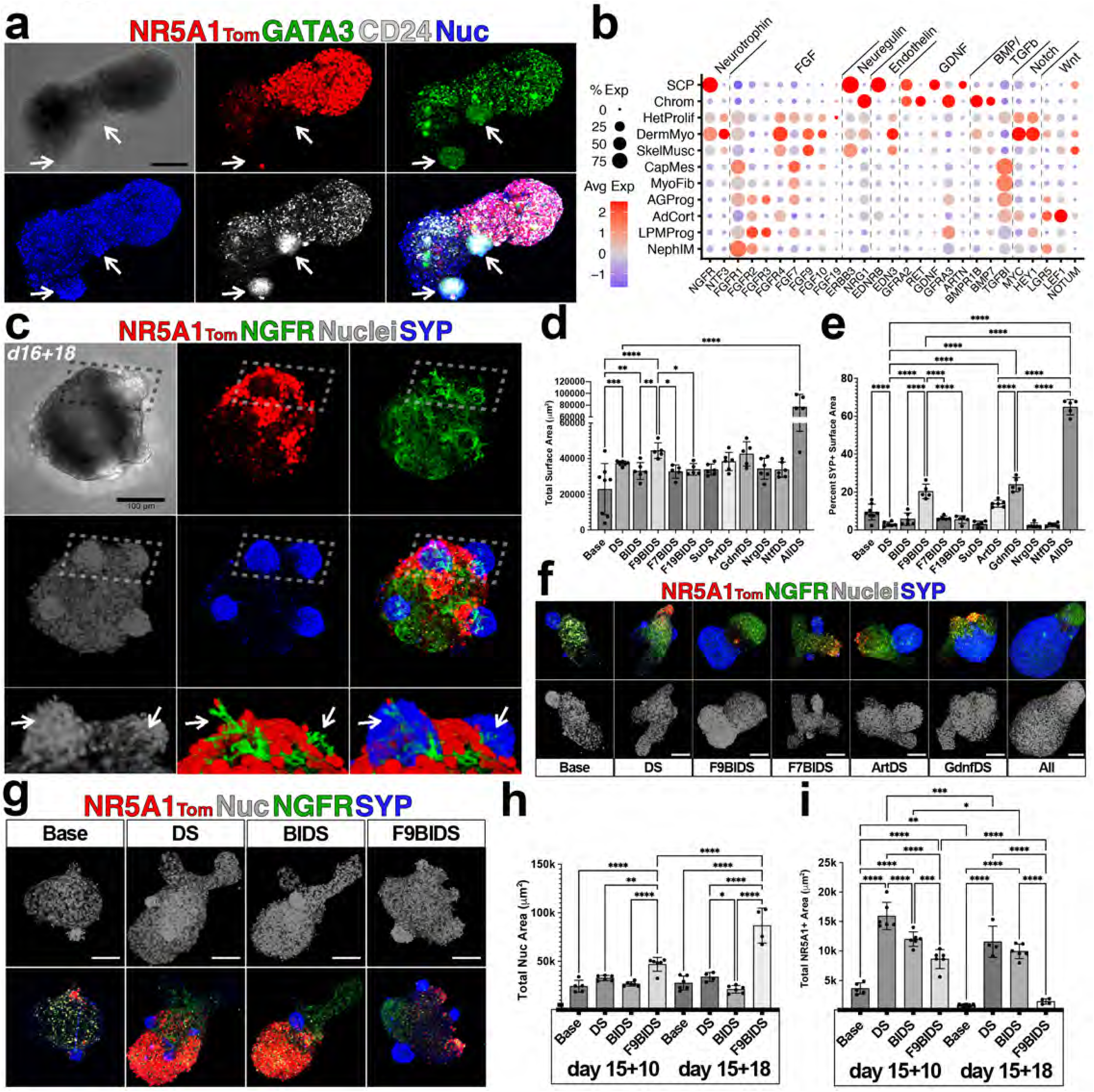
Exogenous FGF9 and GDNF signal stimulus specifically augments chromaffin cell differentiation. a) Immunolabeled with antibodies specific for GATA3 and CD24 highlights coincidence of chromaffin cell clusters (GATA3**^+^**CD24**^+^**) and NT**^+^** cells in individual AGPLOs. b) Dotplot highlighting transcriptional levels of ligand/receptor pairs among AGPLO-derived cells. c) Representative AGPLO following 18 days of growth and containing chromaffin clusters (SYP**^+^**) that are closely associated with NGFR**^+^**(putative SCP) cells; lower panels show volumetric rendering of stroke box. d-f) Total nuclear (d) and SYP**^+^** (e) surface area in NT**^Dim^**organoids cultured for 14 days in bioactive small molecules and/or cytokines, as described in Materials and Methods; representative micrographs of NT**^Dim^** organoids are shown in (f). g-i) Total nuclear (h) and NR5A1+ surface area in NT**^Br^** organoids cultured in Base, DS, BIDS, and FBIDS conditions at 10 and 18 days; representative NT**^Br^**organoids are shown in (g). Scale bars – 100 μm. Error bars in (d, e, h, and i) show standard deviation of the number of replicates represented by dots. The p-values shown are: *p < 0.05, **p < 0.01, ***p < 0.001, and ****p < 0.0001.

To ascertain the influence of each signaling pathway on the cellular composition of AGPLOs, we generated and separately cultured NT**^Dim^**and NT**^Br^** organoids for 14 days in variable combinations of small molecules and recombinant cytokines (Fig 3d-m). Specifically, we tested the activity of fibroblast growth factor 7 (FGF7), FGF9, fibroblast growth factor 19 (FGF19), Su4502 (pan FGF receptor inhibitor), neuregulin 1, neurotrophin 3, Artemin, and glial cell line-derived neurotrophic factor (GDNF). Drawing from previous work^13^ and owing to evident reduction of Notch and TGFb signal activation in SCP and chromaffin populations (Fig 3b), we included DAPT and SB431542 in all conditions excepting baseline control (Base). Additionally, based on potentiation of bone morphogenic protein (BMP)-driven signaling and reduced Wnt-driven signaling in Chrom cells (Fig 3b), we included BMP7 and EndoIWR1 with FGF ligands. SYP**^+^** chromaffin bundles were observed in all conditions, however, Artemin, GDNF, and FGF9 (with BMP7 and EndoIWR) specifically increased the proportion of SYP**^+^** cells (Fig 3d-f and Supp Fig 3a). This increase was observed in the F9BIDS condition within 6 days of organoid culture (Supp Fig 3b-c), however global inhibition of FGF signaling via SU4501 did not significantly reduce total surface area or the proportion of SYP**^+^**cells relative to DS alone (Fig 3d-e and Supp Fig 3a). While FGF9 (relative to cultures including BMP7, EndoIWR, DAPT, and SB431542 alone) was uniquely supportive of increased chromaffin cell volume, it had the converse effect on the proportion of NT**^+^** cells in NT**^Br^**organoids (Fig 3g-i and Supp Fig 3d). Together, these findings identify FGF9, either alone or in combination with known supporting factors in the GDNF family^16^, as a modulator of chromaffin cell differentiation and/or growth.

### FGF9 enriches for bridge sympathoblast phenotype and immature chromaffin cells

To identify mechanisms underlying the influence of FGF9 on shaping the sympathoadrenal compartment, we focused on an earlier timepoint and on 4 culture conditions that consistently enriched for adrenocortical or sympathoadrenal fate (Fig 3g-i). We cultured AGPLOs for 8 days in Base medium (Base), DAPT/SB431542 (DS), BMP7/EndoIWR/DAPT/SB431542 (BIDS), and FGF9/BMP7/EndoIWR1/DAPT/SB431542 (F9BIDS), followed by enzymatic dissociation and scRNASeq (Fig. 4 and Supp Fig 4). Combining these datasets with freshly isolated NT**^Dim^**/NT**^Br^**(***d0 Dim***/***d0 Br***) and d21 AGPLOs (***d21 Pool***) (Fig 2), we performed QC and consolidated Seurat clusters (Supp Fig 4a-d) to assign 15 unique sub-populations (Fig 4a-d). These populations collectively recapitulated principal cellular compartments of the developing AGP filed: adrenogonadal mesodermal derivatives (adrenocortical [**AdCort**], adrenogonadal progenitor [**AGProg**], capsular mesenchyme [**CapMes**], gonadal progenitor [**GonProg**]), neural crest-derived sympathoadrenal cells (Schwann cell precursors [**SCP**], sympathoblasts [**Symp**], chromaffin [**Chrom**]), paraxial mesoderm-derived cells (dermomyotome [**DermMyo**], skeletal muscle [**SkelMusc**]), off-target intermediate mesoderm-derived cells (podocytes [**Podo**]), and lateral plate mesodermal cells (LPM progenitors [**LPMProg**], LPM-derived coelomic mesothelium [**CoelMeso**], LPM-derived mesenchymal progenitors [**LPMMesen**], and LPM-derived stroma [**LPMStrom**]). To provide temporal context for the distribution of each population, we quantified the percentage of total cells of each phenotype (Supp Fig 4e), revealing expansion of the **CapMes**, **DermMyo**, **SkelMusc**, **Chrom**, and **AdCort** populations and reduction of the **GonProg** and **AGProg** populations. Among LPM-derived cells, pseudotime analysis (Supp Fig 4f) suggested differentiation from **LPMProg** to **CoelMeso**, **LPMMesen**, and **LPMStrom** fates, which aligned with population distribution analysis (Supp Fig 4e) showing near-complete absence of LPM-derivatives by day 21, with an initial increase in **CoelMeso**, **LPMMesen**, and **LPMStrom** fates at the expense of **LPMProg** after 8 days.

**Figure 4.**
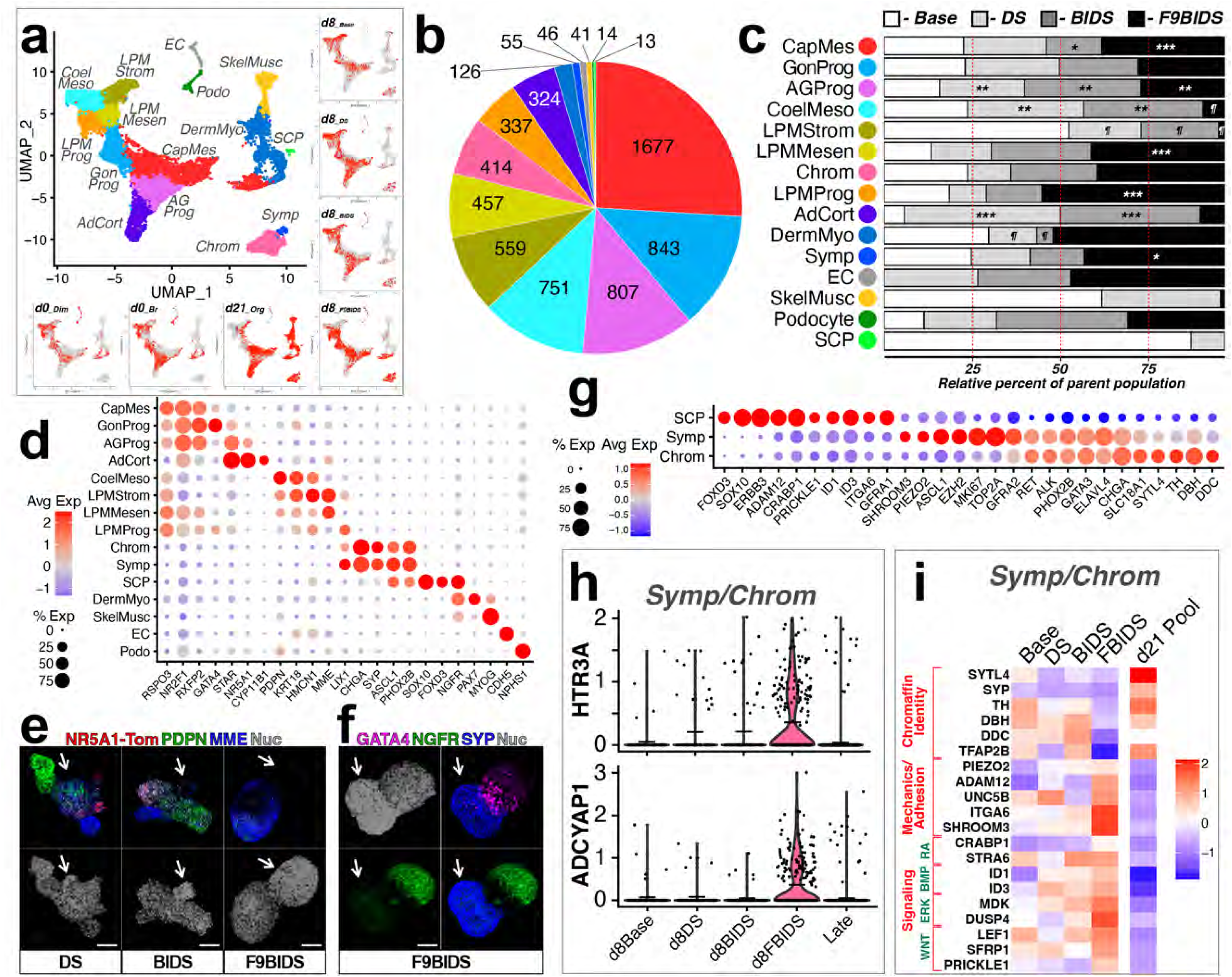
FGF9 expands a bridge sympathoblast and immature chromaffin cell phenotype. a-d) Incorporation of single-cell transcriptomic libraries from AGPLOs cultured for 8 days in either Base, DS, BIDS, or FBIDS conditions. UMAP plots (a) and global (b) versus individualized (c) library contributions of annotated cell populations from each library preparation; asterisks and section signs in (c) denote significantly increased and decreased population levels, respectively. d-f) Dotplot (d) showing unique marker expression among AGPLO-derived populations and immunolabeling (e-f) specific for PDPN, MME, NGFR, GATA, and SYP delineates shift in cell-types represented in various day 8 conditions. g) Dotplot highlighting shifting marker expression as cells progress from SCP ◊ Sympathoblast ◊ Chromaffin fate. h) Violin plot showing expression of HTR3A and ADCYAP1 within the sympathoblast and chromaffin populations cultured in different conditions. i) Heatmap indicating relative expression of transcripts shown between day 8 groups; transcripts are grouped to highlight chromaffin maturation state, mechanosensory/migratory state, and morphogen signal handling. Scale bars – 100 μm.

Focusing on the relative contribution of day 8 growth conditions to each cell type, significant shifts in distribution were elicited in response to FGF9 (Fig. 4c-d): **CapMes**, **LPMMesen**, **LPMProg**, and **Chrom** populations were relatively enriched in the presence of FGF9 at the expense of **CoelMeso** and **LPMStrom** populations; of note, **SCP**s were not observed at all in either the BIDS or F9BIDS condition (Fig. 4c). Immunolabeling of NT**^Dim^**organoids cultured in DS, BIDS, and F9BIDS conditions revealed marker distribution that was consistent with the shift in fates exhibited by transcriptomic analysis (Fig. 4e-f); specifically, localization of podoplanin (PDPN), neprilysin (MME), and GATA4 showed retention of **LPMMesen** (MME**^+^**PDPN**^neg^**) and **GonProg** (GATA4**^+^**) identities at the expense of **CoelMeso** (MME**^+^**PDPN**^+^**) and **LPMStrom** (MME**^+^**PDPN**^dim^**). Sub-analysis of the combined Symp/Chrom population (Fig 4h-i) revealed significant enrichment in the F9BIDS condition of the chromaffin modulator ADCYAP1 and the serotonin receptor gene HTR3A that specifically marks a “bridge-cell” serving as a proliferative intermediate between sympathoblast and chromaffin cell fate^2,16,17^. Additionally, Symp/Chrom cells in the F9BIDS condition exhibited reduced expression of catecholamine enzymatic machinery (TH, DBH, DDC) and increased expression of mechanosensory/migratory and retinoic acid, BMP, ERK, and canonical/non-canonical WNT signaling (Fig 4i), suggesting a relatively immature phenotype^2,17^.

### Sympathetic neuroblast and chromaffin-like cells arise from SOX10 via GDNF-activated proneural exit phase

To clearly define the origin of sympathoadrenal cells in AGPLOs we utilized the SOX10-GFP knock-in hPSC reporter line^18^, which has previously been employed in hPSC differentiation protocols targeting NCC^18^ and sympathoblast/chromaffin^16^ derivatives. Following a differentiation protocol identical to that used for NT line (Fig 1a), FACS-isolated SOX10-GFP**^+^** (SG**^+^**) and NT**^Br^**cells (Fig 5b and Supp Fig 5a-b) were mixed in organoids at ratios of 1:1 (Fig 5c) or combined after two days of individual organoid culture (Supp Fig 5c). In both configurations compound organoids cultured in the presence of DAPT, SB431542, and GDNF (DSG) were comprised mostly of SYP**^+^** cells after 2 weeks of culture (Fig. 5c and Supp Fig 5c). Following optimization of NT**^Dim^** to SG**^+^** cell ratios, NT**^Dim^**/SG**^+^** cells were mixed at 100:1 and generated organoids with >3-fold increase in the proportion of SYP**^+^**surface area (Fig 5d-e), with the SG**^+^**cell origin of chromaffin-like clusters confirmed by residual levels of GFP positivity.

**Figure 5.**
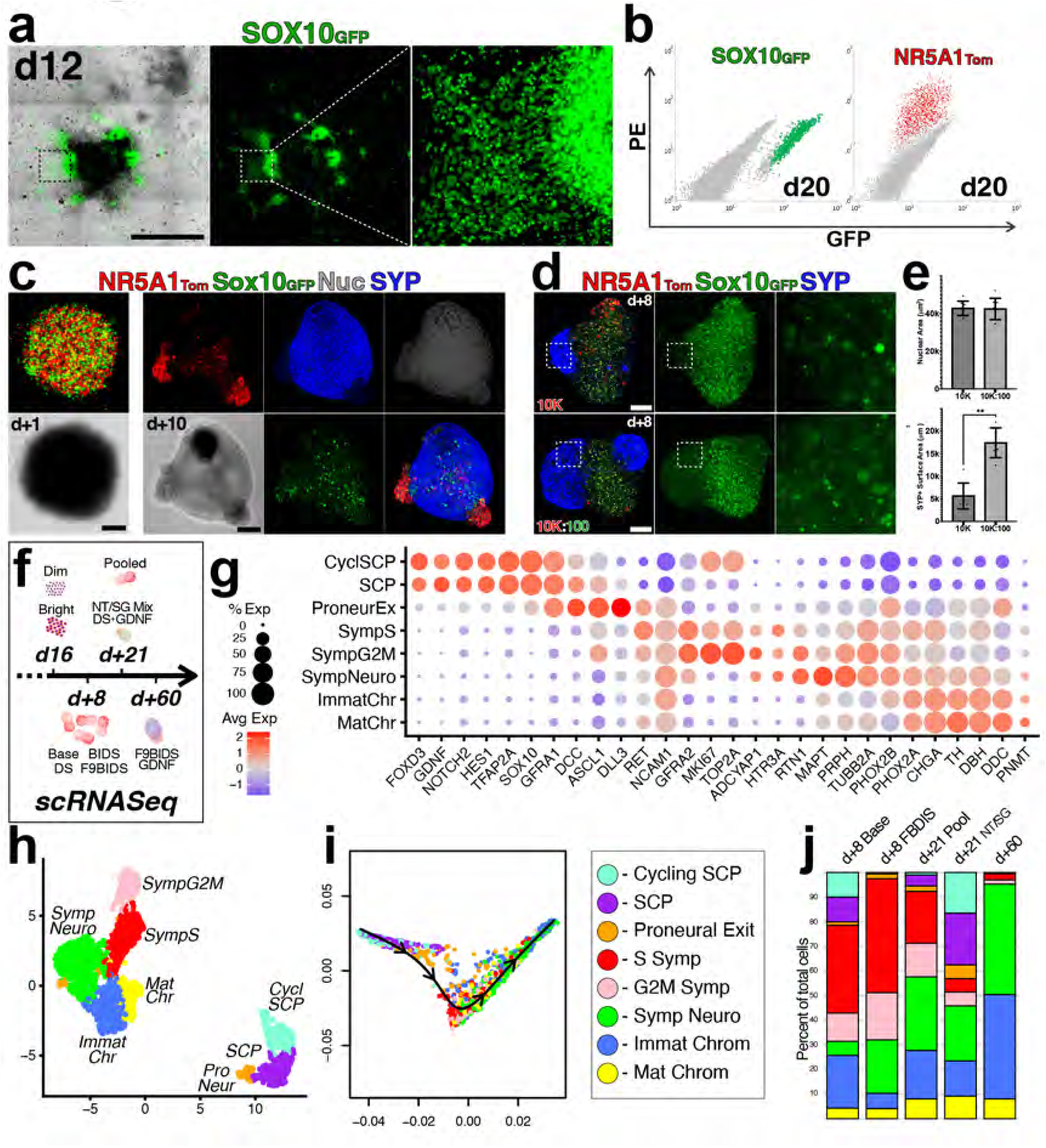
Chromaffin cells derived from SOX10+ SCPs and undergo sympathoadrenal differentiation via a proneural exit state. a) Differentiation of SG and NT hPSCs in identical growth conditions (Fig 1a) results in coincident emergence of NT**^+^** and SG**^+^** derivatives. b-c) FACS sorting (b) and combination of NT**^+^** and SG**^+^**cells at an even ratio (c) result in outgrowth of organoids that are predominantly comprised of SYP**^+^** cells. (d-e) Combination of NT**^+^** and SG**^+^** cells at a 100:1 ratio and growth for 8 days (d) results in significantly increased outgrowth of SYP**^+^**cell area (e). f) Incorporation of two additional single-cell transcriptomic libraries comprised of day 21 mixed SG/NT compound organoids and day 60 NT**^Dim^**-derived chromaffin enriched organoids. g) Dotplot of isolated sympathoadrenal cells from the aggregate library that were re-annotated according to phenotypic placement in the sympathoadrenal differentiation arc. h-j) UMAP plot (h), pseudotime trajectory analysis (i), and quantification of populations within the sympathoadrenal differentiation arc. Scale bars – 100 μm. Error bars in (e) represent standard deviation between 6 replicates. The p-values shown in (e) are **p < 0.01.

To increase the number and/or diversity of the sympathoadrenal lineage available for scRNASeq analysis, we incorporated two additional libraries into the existing dataset (Fig 5f): a group of NT**^Br^**/SG**^+^** mixed organoids (10:1 ratio) that were cultured in the DSG condition for 21 days (*d+21 NT/SG);* and a group comprised of a single NT**^Dim^**-derived organoid that was cultured in F9BIDS + GDNF for 3 weeks and then retained in the DS condition for an additional 39 days (*d+60 F9BIDS/GDNF*). Following QC and isolation of the sympathoadrenal subset based on PHOX2B-expression, a marker of autonomic neuronal lineage (Supp Fig 5e-g), eight clusters encompassing distinct phases along the sympathoadrenal differentiation arc were evident (Fig 5g-i). Based on canonical markers (Fig 5g) and pseudotime analysis (Fig 5i), populations were annotated and ordered as: cycling SCPs (**CyclSCP**) → SCPs (**SCP**) → proneural exit cell (**ProNeur**) → S-phase sympathoblast (**SympS**) → G2M-phase sympathoblast (**SympG2M**) → sympathetic neuroblast (**SympNeuro**) → immature chromaffin cell (**ImmatChr**) → mature chromaffin cell (**MatChr**). Importantly, **ProNeur** cells exhibited a stark transcriptomic switch relative to SCPs that was marked by increased expression of the master proneural transcription factor Achaete-scute homolog 1 (ASCL1)^19,20^, the autocrine Notch inhibitory ligand delta-like ligand3 (DLL3)^21,22^, and the GDNF co-receptor rearranged during transfection (RET)^23,24^, along with decreased expression of the master neural crest cell transcription factor SRY-box transcription factor 10 (SOX10) and the Notch signaling effectors Notch receptor 2 (NOTCH2) and Hes family bHLH transcription factor 1 (HES1). At the population level (Fig 5j), the phenotypic distribution shifted from a predominantly SCP/Symp constitution in d8 organoids to increasing SympNeuro and ImmatChr/MatChr proportions. Combined with the documented role of Notch signal inhibition in SCP escape from lateral inhibition^25,26^ and the observed capacity for exogenous GDNF to expand SYP**^+^** derivatives in NT**^Dim^** organoids (Fig 3d-f), these data delineate a differentiation pathway whereby potentiation of GDNF signaling and autocrine inhibition of Notch mediates sympathoadrenal differentiation of SCPs. Importantly, NCAM1, a known co-receptor for GFRA1 in the absence of RET^27^, is relatively downregulated, but expressed, in SCP populations (Fig 5g and Supp Fig 5h), suggesting a capacity for these cells to respond to GDNF even before acquiring the proneural exit phenotype.

### Combination of NT^Br^ cells with chromaffin-enriched organoids yields ACTH-responsive corticomedullar-like assembloids that generate androgens, glucocorticoids and catecholamines

To reconcile distinct differentiation and growth requirements for generation of medullary (chromaffin) and cortical cells from AGPLOs, we employed a two-stage protocol that first enriched for differentiated chromaffin clusters in NT**^Dim^**/SG**^+^** compound organoids (100:1 cell ratio) and then encased chromaffin clusters with NT**^Br^**cells (Fig 6a). Importantly, encasing freshly generated SG**^+^**-derived organoids or mixing SG**^+^** cells with NT**^Br^** cells at a ratio greater than 1:20 resulted in persistence and hyperproliferation of the SG**^+^** population (Supp Fig 6a-b). By ten days after combination of chromaffin-enriched AGPLOs with NT**^Br^**cells, the NT**^+^** and chromaffin compartments were arranged in a polarized configuration of steroidogenic and catecholaminergic regions that persisted for at least 6 weeks. Production of DHEA-S, androstenedione, testosterone and cortisol was measurable within two weeks (Fig 6d-g) and after 5 weeks of growth, corticomedullar-like assembloids (CLAs) expressing steroidogenic and catecholaminergic machinery (PNMT, Fig 6h) showed increased output of androgens, cortisol, and epinephrine in response to adrenocorticotropic hormone (ACTH) fragment (Fig 6i). In NT**^Br^**:SG**^+^** (10:1 ratio) compound organoids, both steroidogenesis and epinephrine production was evident in response to ACTH as early at 9 days after generation (Supp Fig 6c-g), however these organoids ultimately showed reduced output and reduced proportion of NT**^+^**cells (Supp Fig 6h).

**Figure 6.**
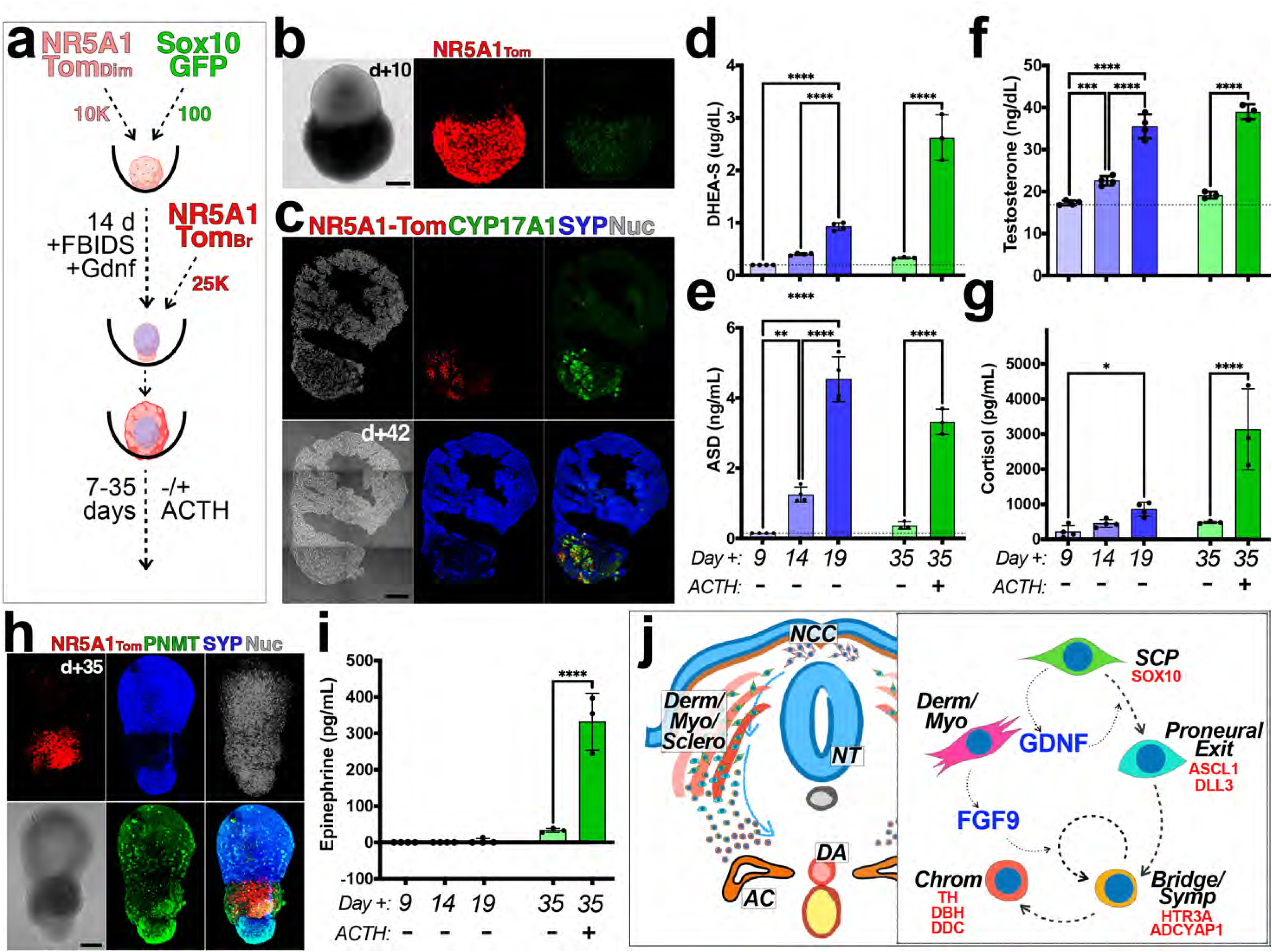
Generation of CMLAs that show integrated cortical and medullary output. a) Schematic for two-step differentiation protocol that combines chromaffin cell-enriched NT**^Dim^**organoids with NT**^Br^** cells to make assembloids. b-c) By 10 days after assembly, cortical and medullary components are polarized, with steroidogenic and catecholaminergic phenotypes retained for at least 6 weeks (c). d-g) Measurement of DHEA-S (d), androstenedione (e), testosterone (f), and cortisol (g) in cell culture supernatant from CMLAs at 9, 14, and 19 days post assembly in the absence of ACTH, and at 35 days in the absence and presence of ACTH. h-i) Expression of PNMT (h) and output of epinephrine (i) from CMLAs cultured in the absence or presence of ACTH. (j) Schematic model describing the relative contributions of GDNF and FGF9-driven effects along the sympathoadrenal differentiation arc.

## Discussion

Due to the disparate anatomical origins of the cortical and medullary compartments of the adrenal gland, a prerequisite to reconstruction of a hPSC-derived model of adrenogenesis is the integration and optimization of protocols that foster the differentiation and combined growth of each component. In this study, we developed an approach that inverts the typical logic of directed differentiation approaches by reconstituting the multi-lineage cell field from which the adrenal primordium emerges and thereby allowing physiologically relevant paracrine relationships within that field to direct fate. We utilized this system to elucidate a novel FGF9-driven signaling mechanism for expanding sympathoadrenal derivatives (Fig 6j), and to generate CMLAs that are responsive to ACTH and capable of steroidogenesis and catecholamine synthesis. This platform advances mechanistic understanding of sympathoadrenal fate specification and establishes an accessible model for interrogating adrenal crosstalk in developmental, physiological and pathological contexts.

The unexpected coincidence of neural crest cell-derived SCPs along with paraxial mesodermal derivatives in AGPLOs elucidated a directional signaling that potentially mirrors physiological paracrine cues within the sympathoadrenal corridor. However, the effect of FGF9 is mechanistically distinct from a lineage-instructive signal. In day 8 AGPLOs, FGF9 enriches the proliferative sympathoblast compartment and HTR3A**^+^** bridge cell phenotype while the mature chromaffin fraction is comparatively unchanged. Extended culture inverts that ratio, indicating that increased volume of chromaffin cells in AGPLOs (Fig 3) results from an expanded bridge rather than a direct effect on chromaffin cells. The transcriptional signature of the F9BIDS sympathoadrenal compartment (catecholamine biosynthetic machinery reduced while a mechanosensory, adhesion and morphogen-handling program is induced) also suggests a larger proportion of cells that have more recently exited the bridge state. Given the source of FGF9 in AGPLOs (dermomyotome-like cells) and documented expression of FGF9 in paraxial/intermediate mesoderm in e8.5 mice^28^, the increase in sympathoblast/chromaffin cell yield may reflect a physiological role for dermomyotome-derived FGF9 in the expansion of a transient progenitor pool in advance of terminal differentiation and integration with the adrenocortical primordium.

Enhanced differentiation within the sympathoadrenal lineage enabled resolution of a scarce but reproducible intermediate correlating to the proneural exit state^2,29^. HES1 is a known suppressor of ASCL1^25,30^, and ASCL1 has been shown to drive DLL3 expression in neuroendocrine cells^31,32^, hence the transcriptional logic of the proneural exit state is consistent and self-sustaining: reduction of NOTCH2 and HES1 releases ASCL1 from suppression, which in turn upregulates DLL3 to further antagonize Notch signal activation. The result is a feed-forward loop by which cells escape lateral inhibition, and it offers a mechanistic explanation for why exit from the SCP pool is stochastic and asynchronous rather than coordinated. Superimposed on this is a reciprocal ligand-to-receptor switch whereby SCPs express GDNF and lack RET, while the exit cell reverses both, extinguishing the ligand and acquiring the receptor. Because SCPs retain GFRA1 and a relatively lower, but present, expression of alternative co-receptor NCAM1^27^, GDNF can also act on the progenitor compartment through the RET-independent route. Hence GDNF, particularly at supraphysiological levels when provided exogenously, can potentially drive signal activation and SCP-exit, rather than just sustaining the signal after the exit decision.

The intermediate proneural exit state resolved here may provide a useful model for interrogating the cellular origins of neuroblastoma. Recent single-cell studies have identified sympathoblasts and immature chromaffin cells, rather than mature chromaffin cells or SCPs, as the transcriptional counterparts of neuroblastoma^33–35^; DLL3 is an established surface target in small-cell lung cancer and other high-grade neuroendocrine tumors^36,37^. Hence, the specific expression of DLL3 in the proneural exit state may identify a developmental intermediate that is retained or re-entered to seed or sustain neuroblastoma; a tractable model for interrogating this cell state could be therapeutically informative.

While CMLAs do not recapitulate that native architecture of the adrenal gland, they recapitulate cortico-medullary crosstalk at a functional level. Epinephrine synthesis requires PNMT, which is induced in vivo by locally high glucocorticoid concentrations delivered from cortical sinusoidal drainage^3,4^. Hence epinephrine is produced almost exclusively in adrenal glands that contain a functional cortex^6^. That CMLAs secrete cortisol and epinephrine from the same tissue, in a polarized steroidogenic–catecholaminergic configuration, indicates that the medullary compartment is not merely adjacent to a steroidogenic one but is being instructed by it. The steroid profile is similarly informative about cortical identity. Concurrent output of DHEA-S, androstenedione and testosterone alongside cortisol indicates that both androgenic and glucocorticoid biosynthetic routes are active, corresponding functionally to zona reticularis- and zona fasciculata-like activity^1,38^. This is notable because the fetal zone, which produces DHEA-S as its principal product, is a primate-specific structure absent from rodents^39,40^ ^40^ and therefore poorly served by mouse models.

Primary adrenal insufficiency, secondary to autoimmune adrenalitis or bilateral adrenalectomy and hormone derangements due to congenital adrenal hyperplasia are managed by fixed-dose steroid replacement that cannot reproduce physiologic, circadian, or stress-responsive secretion^41,42^. A transplantable graft that senses ACTH and secretes on demand would address this shortfall and CMLAs satisfy multiple prerequisites toward this end: they can generate androgens, glucocorticoids and catecholamines; they respond to ACTH; and they maintain a polarized cortico-medullary architecture for at least six weeks. Beyond adrenal cell replacement, the CMLA-platform enables mechanistic and pharmacologic studies that are difficult to perform in rodents, whose medullary architecture, steroid profile and adrenal androgen output differ substantially from human^40,43^. Isogenic introduction of *CYP21A2*, *MC2R*, *MRAP* or *NR0B1* variants into the NT hPSC background would place congenital adrenal hyperplasia, familial glucocorticoid deficiency and adrenal hypoplasia congenita in a human developmental context^44,45^. Finally, because the FGF9 and GDNF manipulations described here allow the sympathoadrenal compartment to be scaled independently of the cortical one, the ratio of the two can be set experimentally, which is prerequisite to using CMLAs either as either disease models or cell-based grafts.

## Limitations

The differentiation hierarchy presented here is inferred from cross-sectional transcriptomes, marker logic, cell-cycle phase and stage distribution rather than from lineage tracing. The proneural exit state is reproducible across independent library analyses but comprises fewer than one hundred cells in each, and both the differentiation trajectory and the proposed Notch-escape mechanism would benefit from clonal barcoding, live reporter imaging, and direct pharmacological manipulation of Notch/GDNF signaling in this system.

Although CMLAs recapitulate the crosstalk between adrenal compartments, they remain incomplete models. We demonstrate responsiveness to ACTH in vitro, but this does not fully represent the complex circuitry of the hypothalamic-pituitary-adrenal axis, and further in vivo studies are needed to assess the competence of these CMLAs as biologic hormone-replacing vehicles. Without vasculature perfusion the current format lacks the counter-current sinusoidal architecture that generates the intramedullary cortisol gradient in vivo^4^ and is therefore unlikely to reproduce the relay with physiological fidelity. Additionally, chromaffin activity in vivo is driven principally by preganglionic cholinergic input rather than by paracrine signals alone^46^, so in the absence of innervation, the secretory dynamics of CMLAs are unlikely to reproduce the kinetics of a stress response.

Recent work from Sasaki lab has demonstrated generation of hPSC-derived organoids that functionally model adrenocortical zonation^47^. Although CMLAs showed functional output, cortical cells were not resolved into glomerulosa, fasciculata and reticularis identities, and aldosterone output was not assessed, leaving mineralocorticoid competence untested. Additionally, the cortical compartment of most CMLAs was relatively smaller than the medullary one, inverting the in vivo proportion. While careful titration of cell inputs can address this, the assembloid configuration of CMLAs is achieved by combination rather than arising by migratory intercalation as it does in the embryo^2,48^.

## Conclusion

By reconstructing the cell field rather than the cell type, we establish a hPSC-based model system in which adrenocortical and chromaffin lineages arise together, occupy compatible axial positions, and signal to one another. We define an FGF9-driven mechanism that expands the proliferative sympathoblast intermediate, and a GDNF/Notch-mediated escape transition through which Schwann cell precursors enter the sympathoadrenal program. Combining these compartments yields assembloids that secrete adrenal androgens, cortisol and epinephrine under ACTH control, thereby providing an accessible model of corticomedullar crosstalk during human adrenogenesis.

## Materials and Methods

### Experimental models

#### Human pluripotent stem cell lines

Two human pluripotent stem cell (hPSC) reporter lines were used in this study. For tracking differentiation within the sympathoadrenal lineage, the previously described SOX10-GFP knock-in hPSC reporter line (SG) was used^18^.Adrenogonadal differentiation and organoid experiments were performed with the H9 human embryonic stem cell line^12^ that has previously been targeted to knock-in green fluorescent protein to the odd-skipped related 1 (OSR1) locus (unpublished). This line was then secondarily targeted to introduce a TdTomato fluorescent cassette into the endogenous *NR5A1* locus to generate the NR5A1-TdTomato reporter line (hereafter NT hPSCs; Supplementary Fig. 1a–b). Briefly, a sgRNA was designed to target a sequence close to the stop codon of the *NR5A1* gene, and the target was cloned into the pX330-U6-Chimeric_BB-CBh-hSpCas9 vector (Addgene plasmid #42230) to make the gene targeting constructs. A donor plasmid containing a 450 bp left homology arm, followed by a P2A-H2B-TdTomato cassette, a floxed puromycin selection cassette, and a 450 bp right homology arm was used as the donor template. The sgRNA and the donor plasmid were electroporated into H9 cells using a Lonza 4D-Nucleofector instrument with *Solution* “Primary Cell P3” and *Pulse Code* “CB-150”. 0.5 μg/ml Puromycin was added to the 3 days post-electroporation for 4 days. Single-cell clones were generated; PCR and sanger-sequencing were used to correctly identify knock-in clones.

hPSCs were maintained under feeder-free conditions on tissue-culture plates coated with hPSC-optimized Matrigel (Corning) in StemFlex Medium (Thermo) at 37 °C under 5% CO2. For passaging or the initiation of differentiation, cells at day 5 post-passaging were dissociated to single cells with Accutase (StemCell Technologies) 6 min at 37 °C, and 10 µM Y-27632 (Tocris) was added for 24 h after replating to enhance survival. Lines were confirmed to be mycoplasma-negative (MycoAlert, Lonza/Cambrex) and were not utilized for more than 8 passages following initial expansion of gene-targeted clones.

## Method details

### Directed differentiation toward NR5A1+ adrenogonadal progenitors

Differentiation toward NR5A1-expressing cells that were used to generate adrenogonadal primordium–like organoids (AGPLOs) was adapted from the previously established floating-aggregate protocol^13^ to support the initial induction phase in suspension conditions followed by a switch to adherent conditions after 3 days (Fig. 1a–b). In brief, hPSCs were differentiated using a staged medium program beginning with 5 days in Induction Medium (comprised of DMEM/F12 with GlutaMAX (Gibco), 2% Fetal Bovine Serum, 100 units/mL penicillin, 100 μg/mL streptomycin, 250 ng/mL amphotericin, and 90 µM 2-mercaptoethanol) followed by Carrying Medium (comprised of DMEM/F12 with GlutaMAX, 10% Knockout Serum Replacement (Gibco), 100 units/mL penicillin, 100 μg/mL streptomycin, 250 ng/mL amphotericin, and 90 µM 2-mercaptoethanol). Embryoid bodies were initially generated by bulk incubation with dispase (1 unit/mL, Gibco) for 15 minutes, collection of floating colonies, two washes in 10x volume of DMEM/F12 with centrifuge at 100x G to collect colonies, and resuspension in Induction Medium including 10 μM Y-27632 (Tocris) and 20 ng/mL BMP7 in Ultra-low Attachment 6-well dishes (3 mL per well). After 24 hours (day 1), 30 μL of 1 mM CHIR99021 was added to each well to bring total concentration of CHIR99021 to 10 μM. After an additional 48 hours (day 3), embryoid bodies from each well were collected and divided into two 10 cm dishes containing 10 mL of Induction Medium including 10 ng/mL BMP7 and 10 μM CHIR99021; 10 cm dishes receiving embryoid bodies for adherent culture were coated with a 250X dilution of Matrigel. Remaining media changes were made using Carrying Medium at day 5: 10 mL with 10 μM CHIR99021, 10 ng/mL BMP7, and 2.5 μM SB431542; day 7: 12 mL with 2 μM CHIR99021, 10 ng/mL BMP7, 25 μM SB431542, 50 ng/mL Sonic hedgehog (SHH), and 100 nM retinoic acid (RA); day 10: 10 mL with 10 ng/mL BMP7, 25 μM SB431542, 25 ng/mL Sonic hedgehog (SHH), 10 μM Endo-IWR, and 10 μM DAPT; day 12: 10 mL with 10 ng/mL BMP7, 25 μM SB431542, 25 ng/mL Sonic hedgehog (SHH), 10 μM Endo-IWR, and 10 μM DAPT; day 16 and successive media changes every third day: 25 ng/mL Sonic hedgehog (SHH), 1 μM Endo-IWR, and 5 μM DAPT.

### Generation of NT^Dim^/NT^Br^ compound adrenogonadal primordium-like organoids (AGPLOs)

Primary differentiation cultures displayed an approximately one-log continuum of NT-reporter intensity (Fig. 1f); cells were sorted into NT***^Dim^*** and NT***^Br^***fractions and recombined to approximate the NR5A1 gradient of the adrenogonadal anlage. For NT***^Dim^*** and NT***^Br^***organoid formation, 20,000 FACS-sorted cells were deposited in U-bottom low-cell-binding 96-well plates (Nuclon) and cultured at 37 °C under 5% CO2 in Carrying Medium. To reconstitute the spatial organization of the primordium, NT***^Dim^***and NT***^Br^*** organoids were generated separately and recombined into compound organoids after 2 days of individual culture (Fig. 1g). Variable combinations of cytokines/small molecules were utilized for culture of AGPLOs. For pooled samples (Fig 2), AGPLOs were cultured initially for 12 days in: Carrying Medium alone (Pool 1), 20 ng/mL BMP7 (Pool 2), 2 μM CHIR99021 (Pool 3), or 10 ng/mL Noggin (Pool 4); and then split into three different conditions for an additional 9 days: base medium, 2 μM CHIR99021, or 1 ng/mL TGFβ1. The three conditions from each Pool were then combined and dissociated for scRNASeq library prep. For day 8 samples (Fig 4), AGPLOs were cultured in designated combinations of BMP7 (20 mg/mL), FGF9 (10 ng/mL), DAPT (10 μM), SB431542 (12.5 μM), and/or EndoIWR (5 μM). For NT/SG mixed AGPLOs (Fig 5), organoids were cultured for 21 days in the presence of 10 μM DAPT, 12.5 μM SB431542, and 10 ng/mL GDNF. For the day 60 sample, the AGPLO was cultured for 21 days in BMP7 (20 mg/mL), FGF9 (10 ng/mL), GDNF (10 ng/mL), DAPT (10 μM), SB431542 (12.5 μM), and EndoIWR (5 μM) followed by an additional 39 days of culture with DAPT (10 μM) and SB431542 (12.5 μM). For assessment of SYP+ area (Fig 3), AGPLOs were cultured in Carrying Medium alone, or with a combination of 10 μM DAPT, 12.5 μM SB431542, 5 μM EndoIWR, 10 μM SU4502, 20 mg/mL BMP7, 10 ng/mL FGF7, 10 ng/mL FGF9, 10 ng/mL FGF19, 10 ng/mL GDNF, 10 ng/mL Artemin, 10 ng/mL Heregulin β1 (Nrg), and/or 10 ng/mL Neurotrophin 3 (Ntf). For all AGPLO cultures, medium was refreshed every 3 days by withdrawal and replacement of 100 μL of medium.

### Sympathoadrenal lineage-tracing and enrichment

To define the origin of sympathoadrenal cells, SOX10-GFP+ (SG+) cells and NT***^Br^*** cells were differentiated in parallel using the protocol above, FACS-isolated, and combined into organoids either by direct mixing at defined ratios (1:1/10:1 NT***^Br^***:SG**^+^**or 100:1 NT***^Dim^***:SG**^+^**) or by recombination after 2 days of separate culture (Supplementary Fig. 5). Mixed organoids were cultured in DAPT, SB431542, and GDNF to promote sympathoadrenal (SYP+) differentiation and then dissociated enzymatically for scRNASeq library preparation.

### Corticomedullar-like assembloid (CMLA) generation and ACTH stimulation

Corticomedullar-like assembloids were generated using a two-stage protocol (Fig. 6a). First, chromaffin clusters were enriched by culturing NT***^Dim^***/SG**^+^** compound organoids (10,000:100 cell ratio) for 14 days in sympathoadrenal-promoting conditions (20 ng/mL BMP7, 10 ng/mL FGF9, 10 ng/mL GDNF, 10 μM DAPT, 12.5 μM SB431542, and 5 μM EndoIWR). Chromaffin-enriched organoids were then encased with 25,000 freshly FACS-sorted NT***^Br^*** cells and further cultured in a combination of 10 μM DAPT and 12.5 μM SB431542. CMLAs were monitored for as long as 6 weeks. For testing of functional output, CMLAs were stimulated with adrenocorticotropic hormone fragment (1 ng/mL). Media was refreshed every five days with 125 μL of withdrawn culture supernatant retained for ELISA analysis.

### Fluorescence-activated cell sorting (FACS)

Differentiation cultures or AGPLOs were dissociated with Accutase for 25 minutes (cultures) or 30 minutes (AGPLOs) min at 37 °C with intermittent trituration utilized for AGPLOs. Enzymatic dissociation cultures were diluted with an equal volume of Carrying Medium, resuspended in FACS running medium (Carrying Medium + DAPI), and passed through 70-µm strainers. Cells were sorted on a FACSJazz (BD Biosciences) based on NT (TdTomato using the PE filter) and/or SG (GFP) reporter fluorescence. Cells were collected in Carrying Media, centrifuged to pellet, and resuspended in Carrying Medium with designated additions of small molecules/cytokines.

### Live imaging, timelapse, and volumetric microscopy

Timelapse imaging of representative compound organoids was performed over 7 days of culture to track reporter and morphological dynamics (Fig. 1h; Supplementary Video 1). Imaging was performed on a LSM 710META Confocal Microscope (Zeiss), using a live incubation chamber (TokaiHit). For volumetric imaging of chromaffin (SYP**^+^**) clusters and their spatial relationship to NGFR**^+^** putative Schwann cell precursors/dermomyotome (Fig. 3c; Supplementary Video 2), whole organoids were fixed, immunolabeled, and imaged across multiple z-planes. Optical sections were compressed and colocalization masking was performed using the Zen imaging suite (Zeiss) to visualize and quantify volumes.

### Immunofluorescence

Organoids were fixed in 4% paraformaldehyde in PBS for 1 h at room temperature, washed in PBS containing 0.2% Tween-20, and blocked in 1% normal donkey serum in PBS/0.2% Tween-20. For immunolabeling, AGPLOs were incubated with primary antibodies for 2 h at room temperature (or overnight at 4 °C), washed, and incubated with Alexa Fluor–conjugated donkey secondary antibodies and 1 µg/mL DAPI for 90 minutes. Live TdTomato and GFP fluorescence was imaged directly without antibody staining. For cryosections, fixed AGPLOs were cryoprotected in 30% sucrose at 4 °C overnight, embedded in OCT, snap-frozen, and cryosectioned at 10 μm. Sections were blocked and stained similarly to whole mount AGPLOs.

### Steroid hormone and catecholamine quantification

For hormone measurements, culture medium supernatants were collected and stored at 4 °C until analysis. DHEA-S, androstenedione, and testosterone were quantified by automated ELISA using the Roche e801 COBAS Analyzer; cortisol and epinephrine were quantified by manual ELISA (Thermo) according to the manufacturer’s specifications.

### Single-cell RNA-sequencing library preparation

Organoids and FACS-isolated fractions were dissociated to single cells, resuspended in 0.1% BSA/PBS, and counted. Single-cell suspensions were processed using the PipSeq Single Cell 3’ RNA Prep Kit (Illumina) following the manufacturer’s protocol. cDNA amplification and library construction were performed per the manufacturer’s instructions, and libraries were sequenced on an Illumina NovaSeq instrument. The libraries analyzed comprised freshly isolated NT***^Dim^*** and NT***^Br^*** cells at day 16 (d0 Dim/d0 Br), pooled day-21 AGPLOs cultured across the cytokine/small-molecule panel described above (d21 Pool), a day-8 series across the Base, DS, BIDS, and F9BIDS conditions, an NT***^Br^***/SG**^+^**day-21 library (d+21 NT/SG), and a long-term F9BIDS+GDNF→DS library (d+60), together with previously published human fetal adrenogonadal (∼4 pcw) and adrenal (∼5 pcw) reference datasets^9^.

### Read mapping and single-cell preprocessing

Sequencing data were demultiplexed and aligned to the human reference genome (GRCh38) with PipSeeker. UMI count matrices were analyzed in R with Seurat (version 5.5.1). For each sample, cells were retained with 200–7,500 detected genes and <10% mitochondrial content; a per-sample minimum-UMI threshold was additionally set by inspection of ranked UMI plots to exclude low-quality cells. Cell-cycle phase was assigned with Seurat’s CellCycleScoring function (S and G2M scores), consistent with the S.Score, G2M.Score, and Phase annotations retained in the metadata.

### Integration, clustering, and cell-type annotation

Counts were normalized and variance-stabilized with SCTransform, regressing out the S.Score and G2M.Score to minimize cell-cycle–driven structure. Sample layers were integrated with Seurat’s IntegrateLayers function using reciprocal PCA (RPCA); Harmony integration was applied in parallel and yielded concordant cluster configurations. The first 30 dimensions were used for FindNeighbors, FindClusters, and RunUMAP. Poor-quality and off-target populations were removed by iterative rounds of UMAP embedding and subsetting. Human fetal reference cells were used to anchor identity, after which hPSC-derived cells were subset, re-clustered, and annotated based on canonical marker genes. Cluster identities were assigned across adrenogonadal-mesodermal (adrenocortical [AdCort], adrenogonadal progenitor [AGProg], gonadal progenitor [GonProg], capsular mesenchyme [CapMes], myofibroblast [MyoFib], nephrogenic/intermediate-mesoderm [NephIM]), neural-crest–derived sympathoadrenal (Schwann cell precursor [SCP], sympathoblast [Symp], chromaffin [Chrom]), paraxial-mesodermal (dermomyotome [DermMyo], skeletal muscle [SkelMusc]), lateral-plate-mesodermal (LPMProg, coelomic mesothelium [CoelMeso], LPM mesenchyme [LPMMesen], LPM stroma [LPMStrom]), and off-target (podocyte [Podo]) compartments.

### Differential expression, module scoring, and HOX profiling

Differentially expressed genes between clusters or conditions were identified with Seurat’s FindMarkers function using the Wilcoxon rank-sum test; genes with an adjusted p-value < 0.05 were considered significant. HOX-cluster expression was profiled across hPSC-derived populations to assign approximate axial (thoracic) identity (Supplementary Fig. 2f–g).

### Pseudotime analysis

Trajectory structure among lateral-plate-mesodermal and sympathoadrenal populations was inferred by pseudotime analysis with Slingshot (learn_graph), with roots placed at the least-differentiated cluster of each lineage (Supplementary Fig. 4f; Fig. 5i).

## Quantification and statistical analysis

Statistical analyses were performed in GraphPad Prism. Group comparisons used two-way ANOVA with multiple comparisons using Tukey’s method, with significance thresholds of *p < 0.05, **p < 0.01, ***p < 0.001, and ****p < 0.0001. The number of biological replicates used is indicated in the corresponding figure legends. Imaging and flow-cytometry results are representative of at least two independent experiments unless otherwise noted.

## Data and code availability

Single-cell RNA-sequencing data and R script have been deposited at GSE344720 and are publicly available as of the date of publication. Any additional information required to reanalyze the data reported in this paper is available from the lead contact upon request.

## Figure Legends

**Supplementary Figure 1.**
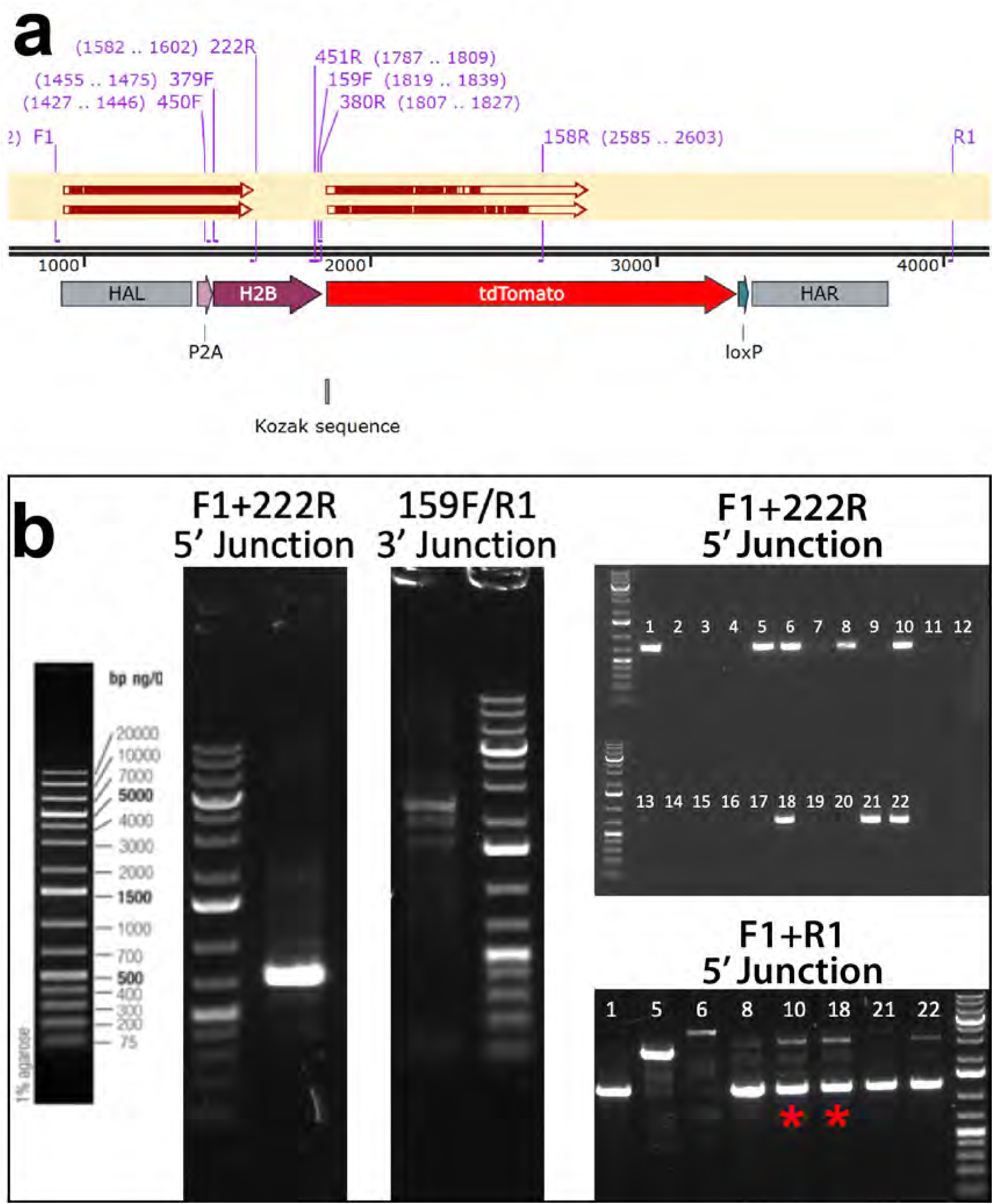
Targeting H9 hPSC line to generate NT hPSCs. a) Schematic diagram of the NR5A1 genomic locus with targeting vector, homology arms (HAL, HAR), and diagnostic sites. b) PCR diagnostic analysis for identification of correctly targeted clones.

**Supplementary Figure 2.**
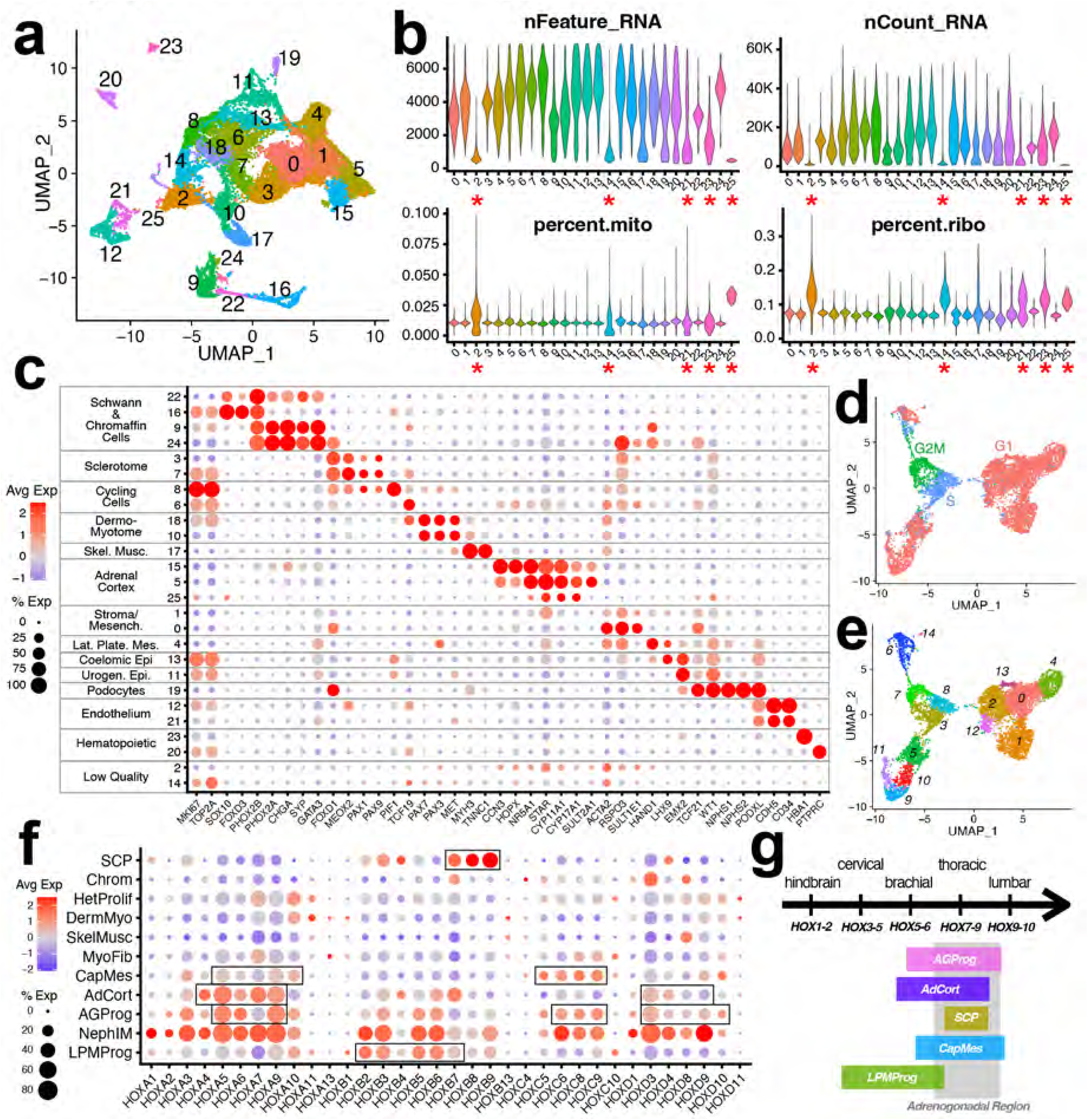
Quality control and axial reference of fetal/AGPLO comparison. a-c) Crude UMAP plot (a), Feature/Count/Mito/Ribo reference (b), and dotplot qualitative assessment of transcriptomic markers (c) from all cells that cleared baseline quality control in the combined d0Dim, d0Br, d21Pool, Week 4 adrenogonadal, and Week 5 adrenal libraries. d-e) Cell-cycle mapping (d) and Seurat clusters (e) of the hPSC-derived fraction of the libraries. f-g) Dotplot (f) and schematic (g) of all HOX genes detected among annotated populations of the hPSC-derived libraries.

**Supplementary Figure 3.**
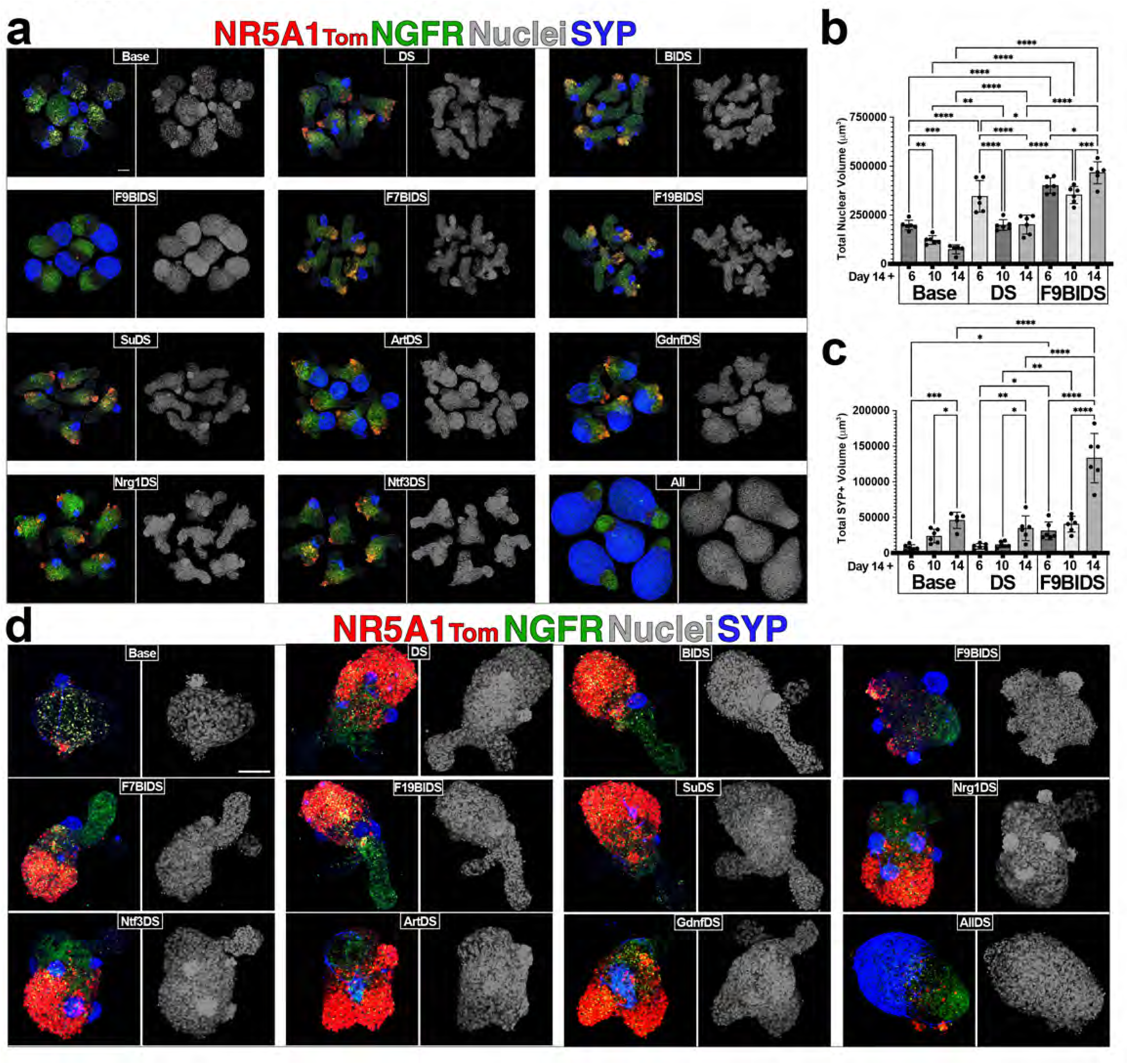
Modulation of chromaffin differentiation in NT^Dim^ and NT^Br^ organoids. a-c) NT**^Dim^** organoids were cultured variable combinations of small bioactive molecules and/or cytokines; Micrographs of organoids after 14 days are shown in (a) and total nuclear (b) and SYP+ (c) volume at 6, 10, and 14 days in either Base, DS, or FBIDS conditions is shown in (b and c). d) NT**^Br^**organoids cultured for 14 days in variable combinations of small bioactive molecules and/or cytokines. Scale bars – 100 μm. Error bars in (b and c) show standard deviation of the number of replicates represented by dots. The p-values shown are: *p < 0.05, **p < 0.01, ***p < 0.001, and ****p < 0.0001.

**Supplementary Figure 4.**
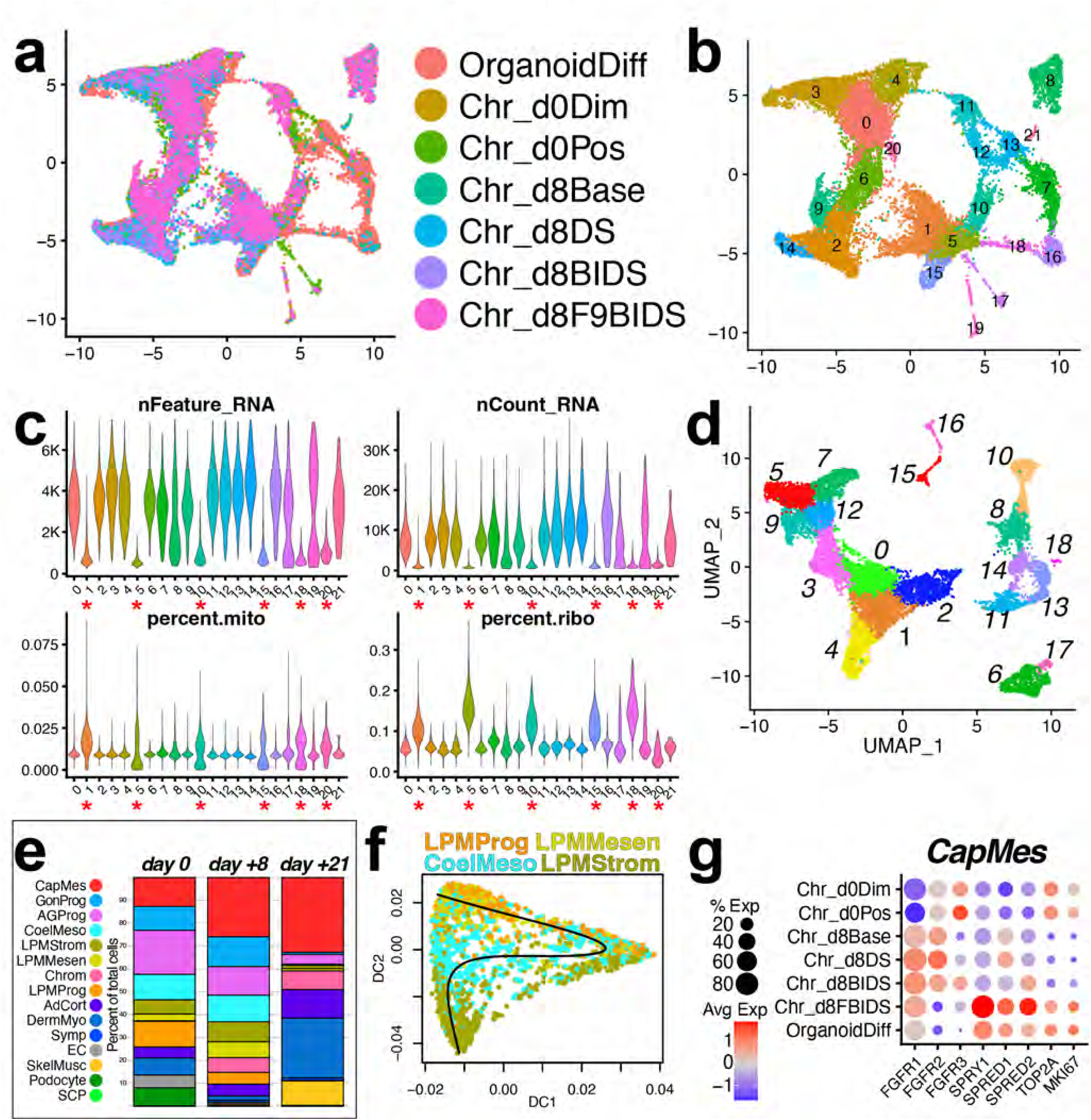
Quality control and subpopulation analysis of day 8 libraries. a-c) UMAP plots of library identities (a), Seurat clusters (b), and Feature/Count/Mito/Ribo reference (c) of the aggregate library containing the combined d0Dim, d0Br, d21Pool, and d8 Base, DS, BIDS, and FBIDS samples. d-e) Seurat cluster map (d) and quantitative breakdown (e) of annotated populations within the aggregate library. f) Pseudotime trajectory map of LPMProg and derivative cells from the aggregate library. g) Dotplot highlighting FGF signaling effectors and proliferative cell markers within the capsular mesenchyme (CapMes) population.

**Supplementary Figure 5.**
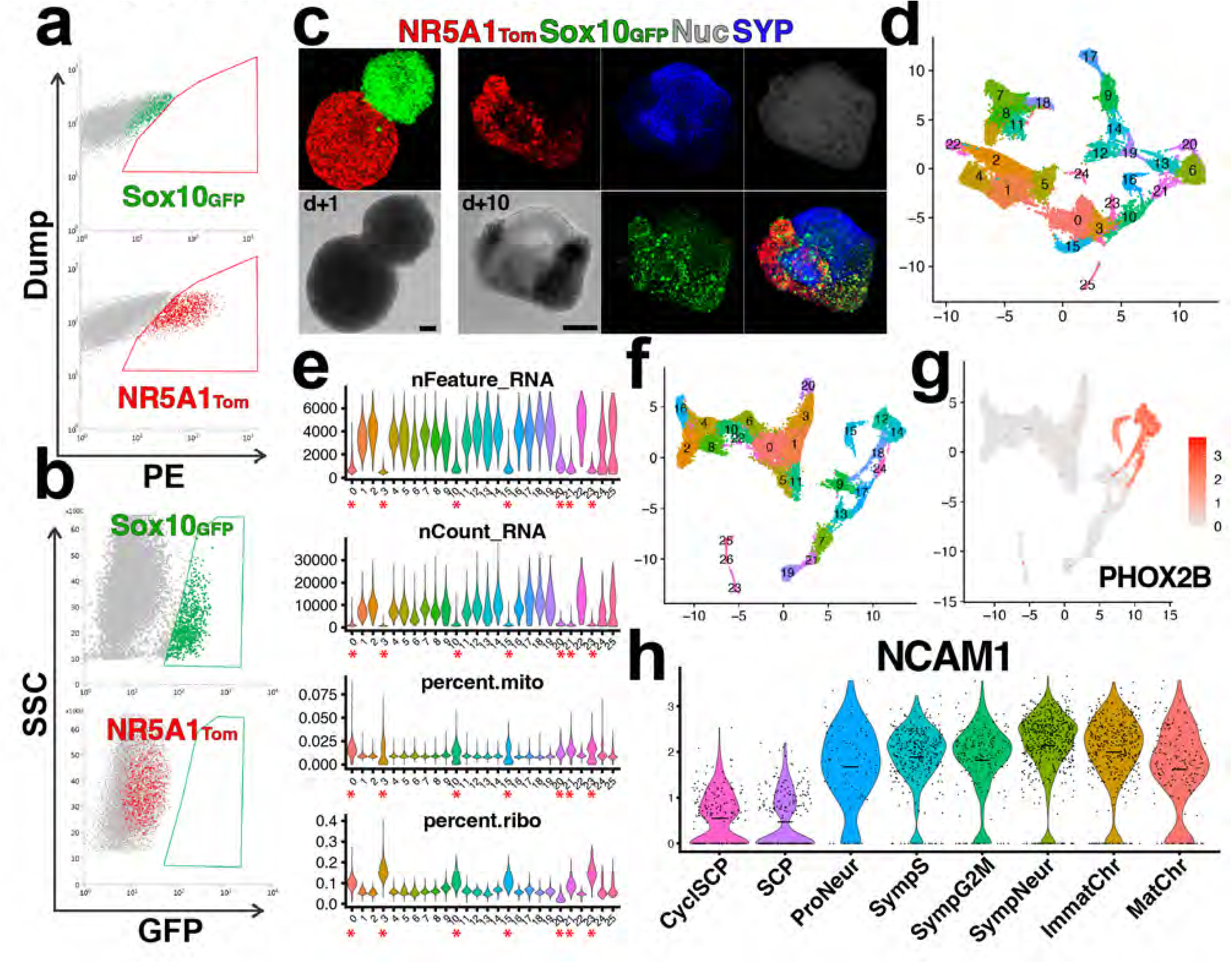
Generation and quality control of sympathoadrenal lineage enriched organoids. a-c) FACS isolation plots of NT**^+^** (a) and SG**^+^** (b) cells and their combination in juxtaposed organoids (c). d-e) Seurat clusters (d), and Feature/Count/Mito/Ribo reference (e) of the aggregate library containing the combined d0Dim, d0Br, d21Pool, and d8 Base, DS, BIDS, and FBIDS, d21 SG/NT mixed, and d60 mature chromaffin samples. f-g) Seurat clusters (f) and PHOX2B feature plot (g) of the post-QC aggregate library. h) Violin plot of NCAM1 expression in the 8 subpopulations within the sympathoadrenal differentiation arc. Scale bars – 100 μm.

**Supplementary Figure 6.**
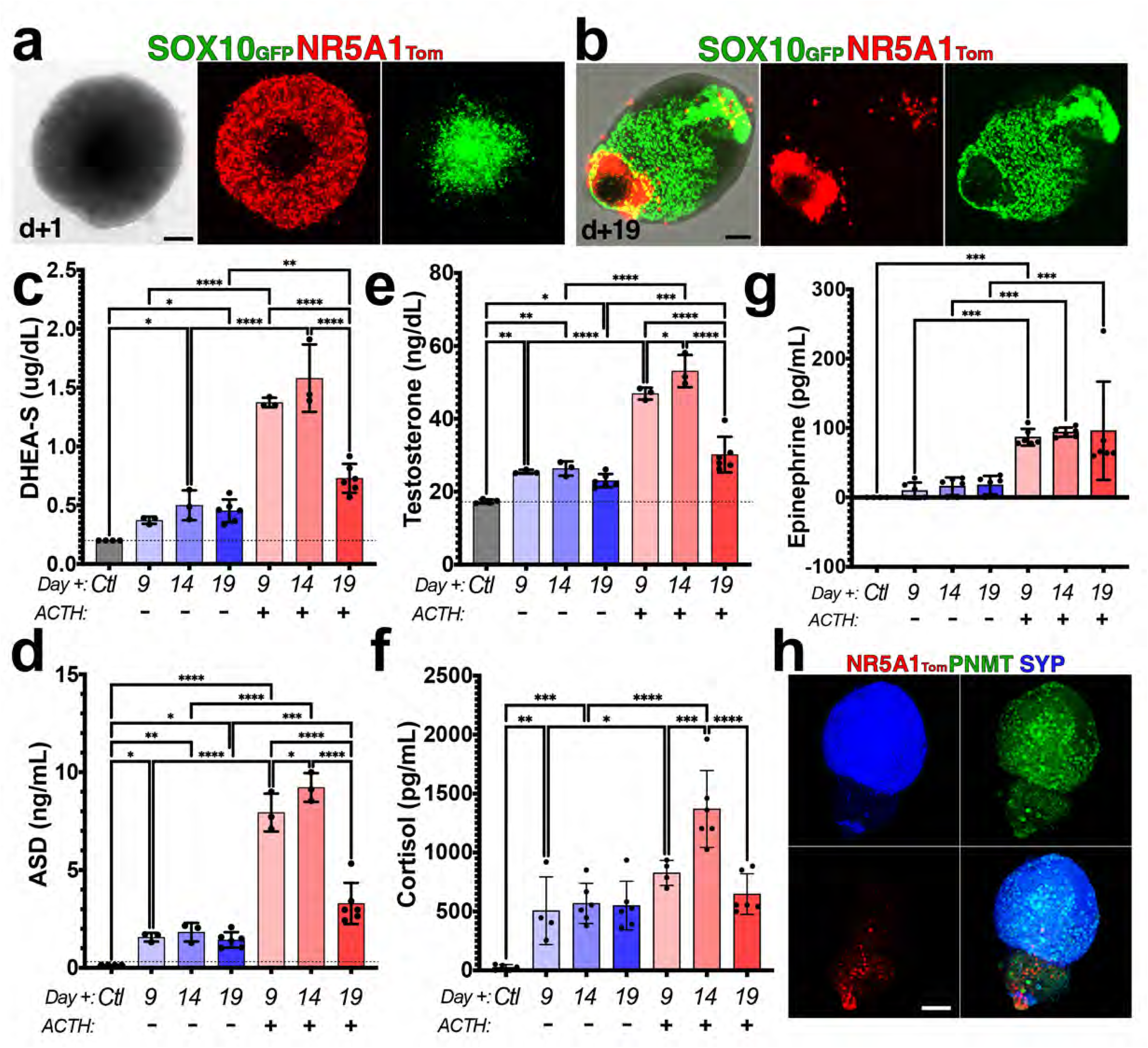
Assembly and functional output of CMLAs. a-b) Assembly of CMLAs by encapsulation of freshly generated SG**^+^** organoids with NT**^Br^**cells (a) results in outgrowths that are enriched for cells that retain high growth potential and SG expression (b). c-g) CMLAs generated from mixing NT**^Br^** and SG**^+^** cells (10:1 ratio) were cultured for 19 days with control or ACTH-containing medium added beginning on day 9. Levels of DHEA-S, androstenedione, testosterone, cortisol, and epinephrine were measured in supernatants collected at 9, 14 and 19 days. h) A representative CMLA at day 35 following chronic stimulation with ACTH beginning at day 9 is labeled with antibodies specific for PNMT and SYP. Scale bars – 100 μm. Error bars in (c-g) show standard deviation of the number of replicates represented by dots. The p-values shown are: *p < 0.05, **p < 0.01, ***p < 0.001, and ****p < 0.0001. The dotted line shows minimum level of detection for the assay.

**Supplementary Table 1. Findallmarkers tables for each aggregate library analysis.**

**Supplementary Video 1. One-week timelapse image capture of AGPLO morphogenesis**.

The video capture one week of differentiation, from two days after isolation from differentiation cultures at day 16 (d16+2) to seven days later (d16+9), captured in 6 minutes intervals. Following capture, the AGPLO was fixed and immunolabeled with antibodies specific for GATA4, CD24, and ALCAM.

**Supplementary Video 2. Volumetric rendering of AGPLO containing chromaffin clusters.**

a) Confocal image capture of a representative AGPLO containing both NT**^+^**, SYP**^+^**, and NGFR**^+^** cell clusters, with three-dimensional rendering illustrating close approximation of SYP**^+^**, and NGFR**^+^** compartments.

## Notes

### Competing Interest Statement

The authors have declared no competing interest.

